# Gene set optimization for cancer transcriptomics using sparse principal component analysis

**DOI:** 10.1101/2025.05.21.655279

**Authors:** H. Robert Frost

## Abstract

A common approach for exploring pathway dysregulation in cancer involves the gene set or pathway analysis of tumor transcriptomic data. Unfortunately, the effectiveness of cancer gene set testing is limited by the fact that most gene set collections model gene activity in normal tissue, which can differ significantly from gene activity found within tumors. To address this challenge, we have developed a bioinformatics approach based on sparse principal component analysis (PCA) for optimizing existing gene set collections to reflect the pattern of gene activity in dysplastic tissue and have used this technique to optimize the Molecular Signatures Database (MSigDB) Hallmark collection for 21 solid human cancers profiled via bulk RNA-seq by The Tumor Genome Atlas (TCGA). Demonstrating the biological utility of our approach, the average survival association of gene set members is improved after optimization for nearly all cancer types and Hallmark gene sets.

## 1 Introduction

Cancer occurs when cells develop a relative growth advantage due to the dysregulation of pathways associated with cell proliferation, survival, or genome maintenance [1]. Understanding the process that leads to pathway alteration, and the impact that pathway dysfunction has on cell and tissue physiology, is therefore critical for an accurate understanding of cancer biology and development of therapeutic interventions. To characterize tumors in terms of biological pathways, researchers have leveraged pathway analysis, or gene set testing, techniques, some specifically optimized for cancer [2, 3], to analyze the large collections of cancer genomic data maintained by projects such as The Cancer Genome Atlas (TCGA) [4].

Although researchers have identified the pathways most frequently impacted by somatic alterations in cancer [1] and progress has been made developing cancer-specific pathway analysis methods that can integrate gene expression and somatic alteration data [2], existing approaches are limited by the misalignment between existing gene set collections and pathway activity within cancerous tissue. Most existing gene set collections, e.g., Gene Ontology [5], represent a tissue-agnostic model of normal gene activity. However, as cataloged by the Human Protein Atlas (HPA) [6] gene expression can differ substantially between different normal tissues and between healthy and malignant samples of the same tissue type [7].

We have previously explored the impact of tissue and cell type-specificity on gene set testing and developed approaches for computing normal tissue-specific [8] and cell type-specific[9] gene set weights, which can be used to select the most biologically relevant gene sets for a particular tissue type or perform p-value weighting after statistical analysis. Although these gene set weights can be used to control for normal tissue or cell type specificity [10], they fail to account for differences between healthy and cancerous tissues. Given the distinct pattern of gene activity found in healthy and cancerous tissue, a gene set testing approach that only accounts for normal tissue-specificity will still generate inflated error rates when applied to tumor expression data. To address the differences between healthy and neoplastic tissue and improve the accuracy of gene set testing on cancer transcriptomics data, we have developed an approach based on sparse principal component analysis (PCA) for customizing existing gene sets to match pattern of gene co-expression in tumors and have used this method to optimize the MSigDB Hallmark collection for 21 solid human cancers profiled by the TCGA. This optimization process and our evaluation approach are detailed in Section 2 with evaluation results in Section 3.

## 2 Data and Methods

### 2.1 Data sources

TCGA data was accessed via the UCSC Xena Datahub [11]. We downloaded and analyzed TPM-normalized RNA-sequencing and overall survival (OS) data for the 21 cancer cohorts listed in Table 1. We applied our gene set optimization method (as detailed in Section 2.2 below) to this cancer gene expression data to optimize the 50 gene sets in version 7.5 of the Molecular Signatures Database (MSigDB) Hallmark collection [12]. To evaluate the effectiveness of this optimization technique, we leveraged gene-level survival association statistics computed for the TCGA data by Uhlen et al. [6]. Specifically, these statistics (*s*_*i,j*_ for gene *i* and cancer type *j*) represent a signed log p-value where the p-value is generated via a Kaplan-Meir (KM) test of the association between the expression of gene *i* and overall survival for TCGA cohort *j* with sign based on the direction of the association (*s*_*i,j*_ *>* 0 for favorable, *s*_*i,j*_ *<* 0 for unfavorable).

**Table 1.**
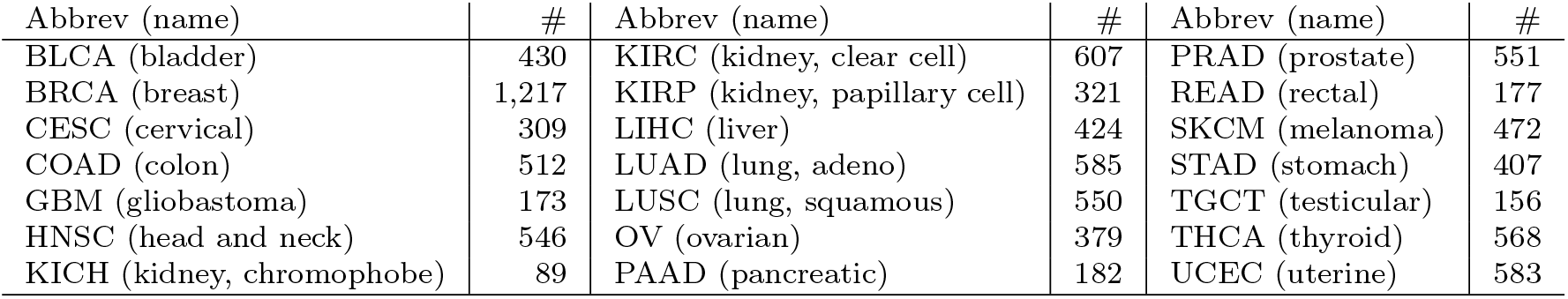
The 21 analyzed TCGA cancer types and number of samples with expression data.

### 2.2 Gene set optimization method

Our approach for cancer-specific gene set optimization leverages the empirical covariance structure of tumor gene expression data. Specifically, our approach seeks to identify one or more subsets of gene set members that are significantly co-expressed in tumors of a specific cancer type. This approach is motivated by the assumption that genes which play a biologically important role in a particular gene set/pathway will be positively co-expressed in the relevant normal tissue or tumor type [13]. To identify the co-expressed groups of genes in each set, we leveraged our Eigenvectors from Eigenvalues Sparse Principal Component Analysis (EESPCA) [14] method. The EESPCA method is based on the recently detailed formula for computing normed, squared eigenvector loadings of a Hermitian matrix from the eigenvalues of the full matrix and associated sub-matrices [15]. Importantly in this context, the sample covariance matrix 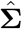 is Hermitian, which implies that squared normed PC loadings (|*v*_1_|^2^ for the first PC) can be computed as a function of PC variances for the full data set and all of the leave-one-out variable subsets. Specifically, the normed squared loading for variable *j* in the first PC can be computed as: 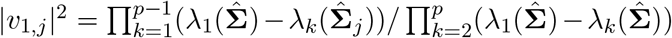, where 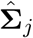 represents the sample covariance sub-matrix with variable *j* removed and *λ*_*k*_(**Σ**) is the *k*^*th*^ eigenvalue of **Σ**. EESPCA is based on an approximate version of this formula, 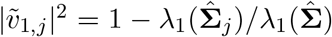, that can be efficiently computed for very large data matrices using the method of power iteration. Importantly, the ratio between the approximate and true eigenvector loadings, 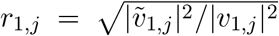, provides information regarding the true sparsity structure of the population PCs and is used by the EESPCA method to generate sparse PC loading vectors. Relative to the state-of-the-art sparse PCA methods of Witten et al. [16], Yuan & Zhang [17] and Tan et al. [18], the EESPCA technique offers a two-orders-of-magnitude improvement in computational speed, does not require estimation of tuning parameters via cross-validation, and can more accurately identify true zero principal component loadings across a range of data matrix sizes and covariance structures.

To optimize membership for a specific gene set and TCGA cohort, the EESPCA method is used to compute the first three sparse PCs using the normalized RNA-seq counts for the set genes. The loadings of these sparse PCs are then used to create three different optimized versions of the target set by removing all genes that have a zero loading on the first PC, the first two PCs or the first three PCs. This approach supports the scenario where a set is comprised by up to three independent co-expressed blocks of genes.

### 2.3 Evaluation

To evaluate our cancer-specific gene set customization approach, we compared the *s*_*i,j*_ survival association statistics for genes in unoptimized and optimized gene sets. If the optimization process is preferentially retaining genes that play an important role in the biology of the target cancer, we expect that the average magnitude of the surival statistics will be higher for the optimized set than for the unoptimized set. If the *m* genes in the original set are split into *m*^*o*^ in the optimized set and *m*^*r*^ that are removed, this expectation can be stated mathematically as 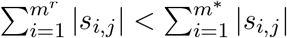. When this inequality holds, we can conclude that the quality of the gene set improved after optimization, otherwise, it worsened. To assess the statistical significance of the change in survival statistics, directional Wilcoxon rank sum tests were performed comparing the survival statistics of retained vs removed genes. If the relevant directional test was marginally significant at *α* = 0.05, the optimized set was deemed to be either ‘significantly improved’ or ‘significantly worse’. Overall, each optimization result was assigned to one of four distinct outcome classes: significantly improved, improved, worse, and significantly worse.

A related evaluation can be performed by comparing the magnitude of the survival statistics for cancer type *j* between a set optimized according to the expression data for cancer *j* (e.g., liver) and the same set optimized for a distinct cancer type *k* (e.g., breast). Our expectation in this case is that the average magnitude of the survival statistics will be larger when optimization was performed using expression data from the same cancer type than from a distinct cancer type, e.g., 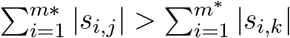.

## 3 Results

### 3.1 Proportion of gene set annotations retained after optimization

Figure 1 visualizes the proportion of gene set annotations retained after optimization for each of the 21 target TCGA cohorts and number of PCs. As expected, the number of genes retained after optimization increases as the number of PCs considered increases. The average proportion retained across all cohorts and Hallmark gene sets was 0.30 for 1 PC, 0.53 for 2 PCs, and 0.65 for 3 PCs. While the impact of optimization is similar for most cancer types, ovarian cancer (OV) is a noticable outlier with almost no genes removed when 2 or 3 PCs are considered, which implies that the higher order sparse PCs computed on the ovarian cancer expression data have very few zero values.

**Figure 1.**
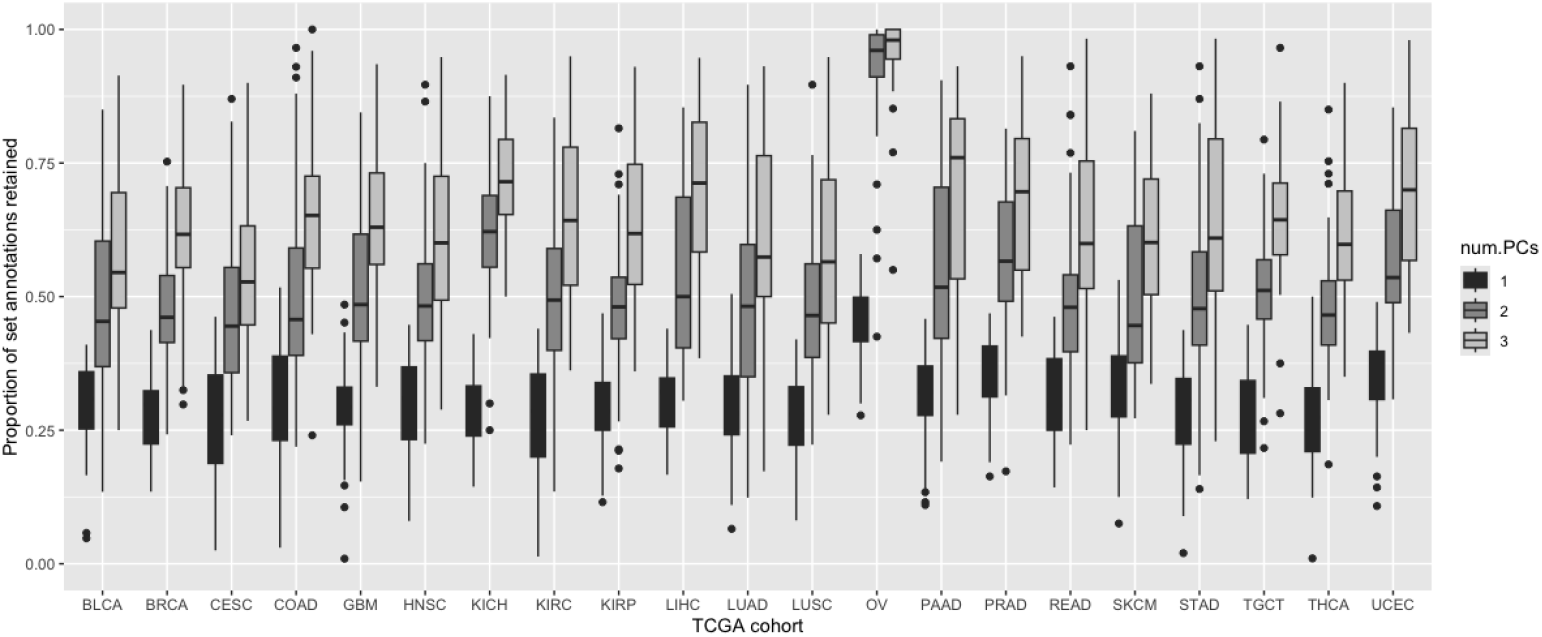
Proportion of Hallmark gene set annotations retained after optimization on the bulk RNA-seq data for each TCGA cohort using 1 to 3 sparse PCs.

### 3.2 Optimization outcomes for each TCGA cohort

Figure 2 visualizes the proportion of gene sets whose optimization results fall into one of the four outcome categories: significantly improved, improved, worse, and significantly worse (see Section 2.3 for details). Averaged across all of the cohorts and PC numbers, the average gene-level survival association significantly improved for 25.3% of the gene sets, improved for an additional 37.9%, worsened for 29.9% and significantly worsened for only 6.9%. Optimization using 3 PCs had the best average outcome (22.8% significantly improved, 44.2% improved, 27.3% worse, and 5.7% significantly worse). Importantly, optimization improves the survival association of annotated genes for well over half of the Hallmark gene sets for almost all of the TCGA cancer types (prostate (PRAD) and testicular (TGCT) cancers being the notable exceptions).

**Figure 2.**
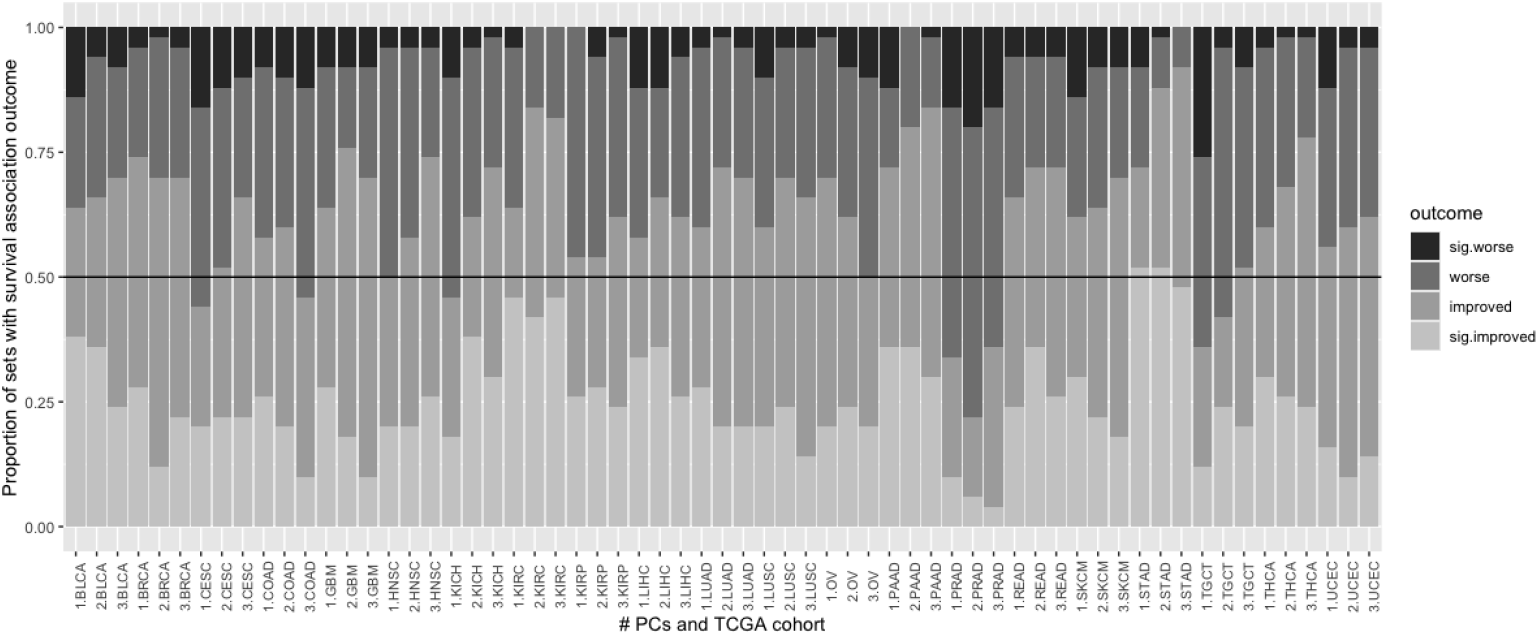
Proportions of the Hallmark gene sets by outcome class after optimization on the bulk RNA-seq data for each TCGA cohort using different numbers of PCs.

Figure 3 provides a similar outcome visualization for the case where gene sets optimized for one TCGA cohort are evaluated using survival statistics from distinct cohorts. As expected, the proportion of improved gene sets is lower in this case as compared to the scenario when survival statistics and expression data are based on the same cohort. Specifically, the average gene-level survival association significantly improved for 17.1% of the gene sets (vs 25.3%), improved for 39.5% (vs 37.9%), worsened for 35.1% (vs 29.9%) and significantly worsened for 8.3% (vs 6.9%). Surprisingly, our optimization method was still generally effective in this case which implies that the biologically relevant gene co-expression pattern is broadly similar across many solid human cancers. One notable outlier is liver cancer (LIHC), which is likely due to the fact that liver tissue has a relatively large number of highly tissue-specific genes [6].

**Figure 3.**
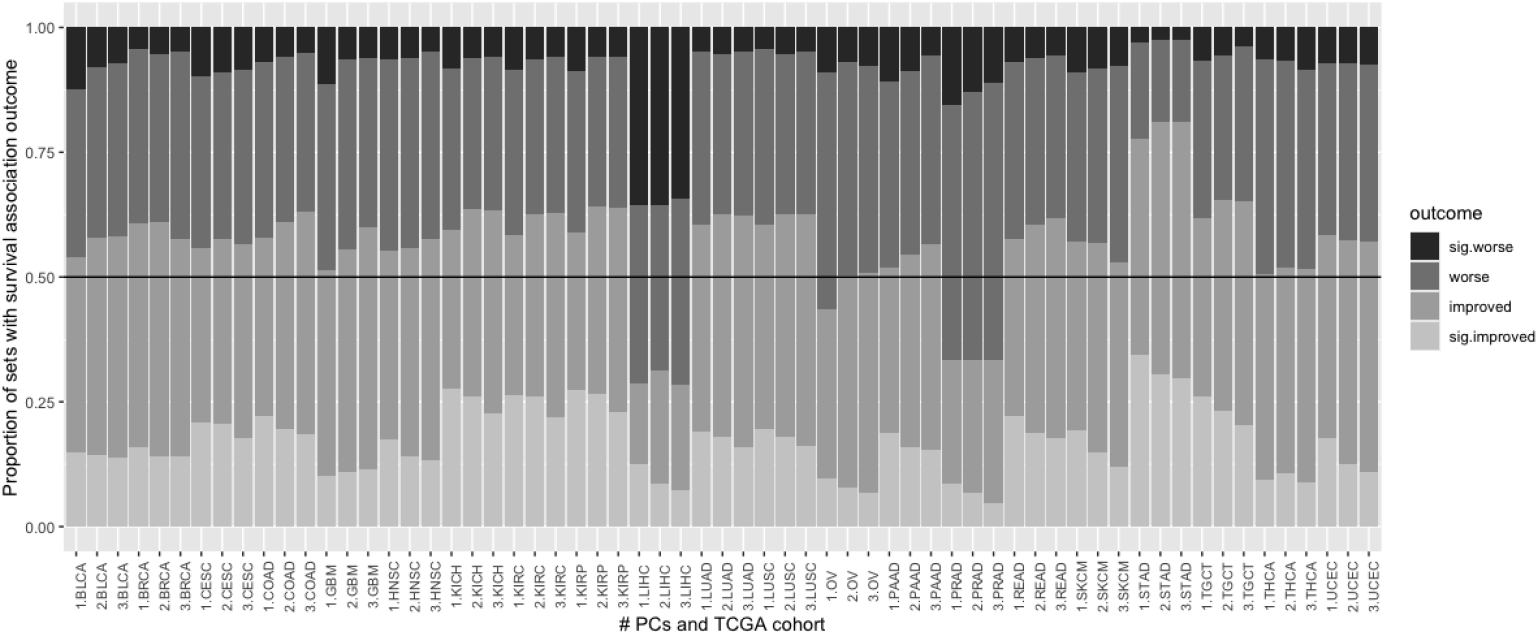
Proportions of the Hallmark gene sets by outcome class after optimization on the bulk RNA-seq data from distinct TCGA cohorts.

## 4 Conclusions and future directions

To address the mismatch between public gene set collections, which are typically based on gene activity within normal tissues/cells, and the pattern of gene expression in cancer, we developed an optimization method based on the sparse PCA of tumor gene expression data. The results in this paper demonstrate that this approach can effectively retain genes whose expression is significantly associated with cancer survival. Importantly, for the optimization results based on 3 PCs for 21 solid cancers and 50 Hallmark gene sets, the mean survival association was improved in 63% of the cases with four times as many sets having a statistically significant increase in survival association strength than a significant decrease. Of course, not all cancer types and gene sets benefit from this optimization and evaluation based on the association with overall survival is only a proxy for general biological relevance. Given the heterogeneity of the results, the practical application of this type of approach would only use the optimized sets on non-TCGA data that saw a significant improvement on the relevant TCGA cancer type.

For the full-length version of this short paper, we plan to 1) extend the evaluation results to all MSigDB collections, 2) create public repository of optimized gene sets, 3) include gene set-specific results in addition to cancer type-specific results, and 4) explore other evaluation metrics (e.g., prognosis prediction using single-sample gene set statistics, recurrence-free survival vs. overall survival, association with non-survival endpoints, etc.).

## Conflict of interests

The authors have no conflicts of interest to declare.

## Acknowledgments

I would like to thank Xingyu Zheng for her help evaluating an early prototype of this approach.

## Funding

This work was funded by National Institutes of Health grants R35GM146586, R21CA253408, P20GM130454 and P30CA023108.

## Notes

### Competing Interest Statement

The authors have declared no competing interest.

